# The knockout effect of low doses of gamma radiation on hepatotoxicity induced by *Echis Coloratus* snake venom in rats

**DOI:** 10.1101/705251

**Authors:** Esraa M. Samy, Esmat A. Shaaban, Sanaa A. Kenawy, Walaa H. Salama, Mai A. Abd El Fattah

## Abstract

*Echis Coloratus* is the most medically important viper in Egypt causing several pathological effects leading to death. Gamma radiation has been used as a venom detoxifying tool in order to extend the lifespan of the immunized animals used in antivenin production process. Thus, the aim of this study is to assess the effects of increasing doses of gamma radiation on *Echis Coloratus* in vivo through biochemical and histological studies. The results revealed a significant increase in the levels of AST, ALT, ALP and glucose of sera collected from the rats injected with native *Echis Coloratus* venom compared with the non-envenomed group. On the other hand, biochemical parameters of sera of rats administrated with either 1.5 kGy or 3 kGy irradiated venom were significantly decrease compared with the native venom envenomed group at 2h, 4h and 24h post envenomation. In addition, these results were confirmed by histological studies of rats’ livers. Correspondingly, the sublethal dose injection of native *Echis Coloratus* venom induced significant alterations in the histological architecture of liver after 2, 4 and 24 h of injection. Concurrently, the administration of both 1.5 kGy and 3 kGy gamma irradiated venom showed fewer histological alterations compared with the native group. In conclusion, the present findings support the idea of using gamma radiation as an effective venom detoxification tool.

## 1. Introduction

*Echis Coloratus* is a highly venomous snake which causes numerous bites in Egypt and results in high levels of morbidity, disability and mortality. It belongs to family *Viperidae* which is one of the largest and most widely distributed vipers present in North Africa and Middle East ***(WHO, 2016)*.**

Envenomation by *Echis Coloratus* is characterized by swelling and pain which are the earliest symptoms appearing within minutes of the bite. In severe envenomation the swelling can extend to whole of the limb and can lead to limb amputation ***(Ali et al., 2004; Vaiyapuri et al., 2013)*.** *Echis Coloratus* venom is vigorously attacked liver causing hepatocellular damage, hemorrhage, elevation in liver enzymes and depletion in glycogen ***(Al-Jammaz, 2003)*.** Venom is composed of a cocktail of biomolecules include enzymes, peptides, carbohydrates, lipids, nucleosides, amines and inorganic metal ions such as sodium, calcium, potassium, magnesium, zinc, iron, cobalt, manganese, and nickel (***Burin et al., 2018***).

Venoms of *Viperidae* are rich sources of protease enzymes which are responsible for digestion of tissue proteins, so venoms rich in proteases produce marked tissue destruction ***(Tan et al., 1989; Braud et al., 2000)*.** Phospholipase A_2_ (PLA_2_) is another important enzyme in *Echis Coloratus* venom; they are small enzymes which hydrolyze phospholipids releasing lysophospholipids and fatty acids. PLA_2_ enzymes interfere in many normal physiological processes such as presynaptic and postsynaptic neurotoxicity, myotoxicity, platelet aggregation initiation, convulsions, hypotension and damage to Liver, kidney, lungs, testis and pituitary gland ***(Middleton et al., 2000; Teixeira et al., 2003)*.**

Venoms are weakly immunogenic and very toxic, causing difficulties for antivenom production. Therefore, the use of detoxified venoms can be very helpful in many aspects ***(Caproni et al., 2009)*** but it is important to be sure that the detoxified venoms do not lose their immunogenicity. Gamma radiation has been shown to be effective for attenuating venom toxicity while maintaining immunogenicity ***(Shaaban, 2003)*.** Since there is a close relationship between the structure and the biological activity of macromolecules, some alteration appeared to be the most possible explanation for the radiation effects. Loss of function of protein by irradiation is not usually due to breaking peptide bonds, or otherwise, disrupting the primary skeletal structure of peptide chain. It may result from a break in the hydrogen or disulfide bonds which in turn can result in a disorganization of the internal relationships of side chain groups, or an exposure of amino-acid groups, resulting in change in biological activity ***(Hayes and Kruger, 2001; Samy et al., 2018)*.**

Thus, the aim of this study was to evaluate the effectiveness of both 1.5 kGy and 3 kGy gamma rays as a detoxification tool of *Echis Coloratus* snake venom by some biochemical analyses and histological examination of liver tissues of experimental rats.

## 2. Materials and methods

### 2.1. Animals

#### 2.1.1. Rats

Male Wistar albino rats (150-180 g) were used for determination of histopathological and biochemical changes after intra-peritoneal injection of venom. Animals were kept under appropriate conditions of temperature, humidity and light and were allowed free access to food and water ad libitum.

#### 2.1.2. Snakes

The crude venom was obtained from *Echis Coloratus* snakes collected from Sinai, Egypt and kept in laboratory unit of Medical Research Centre, Faculty of Medicine, Ain Shams University. The venom was freeze-dried and kept in the refrigerator at 4°C until used.

### 2.2. Irradiated Facilities

(1mg/ ml) of the venom dissolved in saline (0.9% NaCl) was irradiated with a dose level of 1.5 kGy or 3 kGy gamma rays in the National Center of Radiation and Research Technology, Cairo, Egypt, using cobalt-60 gamma cell 220, Atomic Energy of Canady Limited (AECL), Canada, in the presence of O_2_ at room temperature. The radiation dose rate was 1.2 Gy per second.

### 2.3. Methods

#### 2.3.1. Biochemical analyses of native and (1.5 kGy and 3 kGy) gamma irradiated venom

Four groups each subdivided to 3 subgroups, each subgroup contain three rats. Three envenomed groups were intra-peritoneal (i.p.) injected with 0.672 mg/kg of venom; a sublethal dose of LD_50_ (Median lethal dose) determined in a previous study **(*Samy et al., 2015)***. The control group was injected by saline (0.9% NaCl). The sera were collected from the retro-orbital vein of rats after 2h, 4h, and 24h of venom envenomation, and centrifuged at 5000 rpm for 10min. Biochemical analyses of collected sera were performed using UV-Spectrophotometer and specific kits purchased from Biodiagnostic Company, Egypt. The levels of glucose, AST (aspartate aminotransferase), ALT (alanine transaminase) and ALP (alkaline phosphatase) were determined in sera of both the control and envenomed groups.

#### 2.3.2. Histopathological study

The histopathological studies of liver of rats were performed using light microscope. First, rats were sacrificed by decapitation under anesthesia at 2, 4 and 24 h post envenomation, the liver was removed and immediately fixed in 10% formalin solution, kept until become hard enough to be sectioned and embedded in parrafin blocks. Sections of 5 mm thickness were stained with hematoxylin and eosin (H&E) and mounted on slides for examination as described by ***Banchroft et al. (1996)*.**

#### 2.3.3. Statistical analysis

Data were reported as mean ± standard error (S.E) and were subjected to a one-way analysis of variance (ANOVA) followed by Tukey- Kramer multiple comparison test for comparisons between group means. Values of p lower than 0.05 was considered as significant.

## 3. Results

### 3.1. Biochemical Analysis

Compared to the control group, the biochemical analyses of rats’ sera received native venom showed a significant increment in glucose, ALT, AST and ALP levels (p < 0.05) at 2, 4 and 24 h post envenomation. Additionally, the highest peak was noted at 4 h of venom injection. Whereas, the measurements of these parameters in the rats’ sera received 1.5 and 3 kGy gamma irradiated venoms were reduced compared to the native venom treated group. Moreover, the biochemical parameters were decreased to normal levels in rats envenomed with 3 kGy gamma irradiated venom after 24 h post envenomation (Table 1).

**Table (1).**
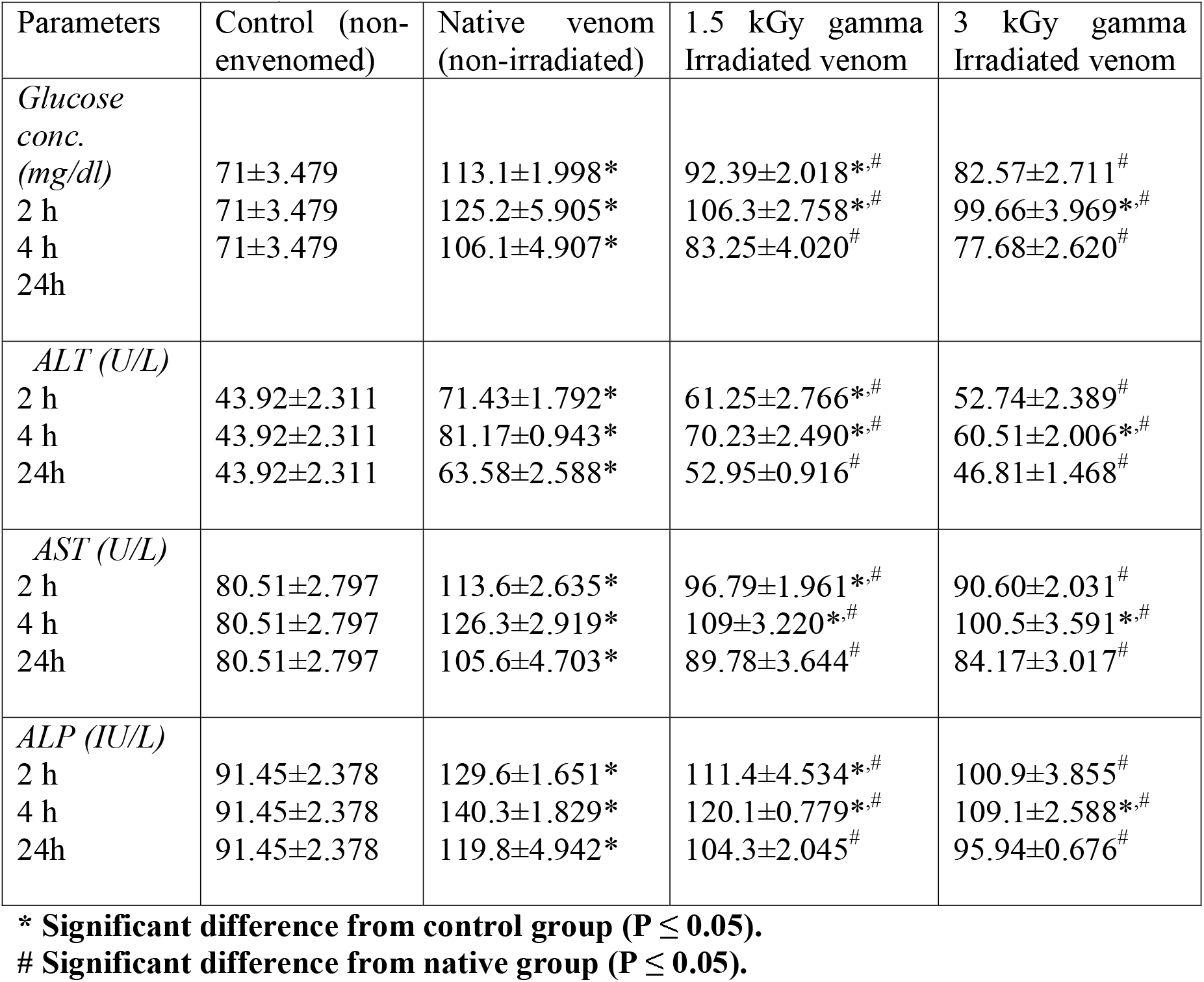
Effect of sublethal dose (0.672mg/kg, i.p.) of native, 1.5 and 3 kGy gamma irradiated venom on glucose, ALT, AST and ALP levels at 2, 4 and 24 h. (Mean ± standard deviation)

| Parameters | Control (non-envenomed) | Native venom (non-irradiated) | 1.5 kGy gamma Irradiated venom | 3 kGy gamma Irradiated venom |
| --- | --- | --- | --- | --- |
| *Glucose conc. (mg/dl)* |  |  |  |  |
| 2 h | 71±3.479 | 113.1±1.998* | 92.39±2.018*.<sup>#</sup> | 82.57±2.711<sup>#</sup> |
| 4 h | 71±3.479 | 125.2±5.905* | 106.3±2.758*.<sup>#</sup> | 99.66±3.969*.<sup>#</sup> |
| 24h | 71±3.479 | 106.1±4.907* | 83.25±4.020<sup>#</sup> | 77.68±2.620<sup>#</sup> |
| *ALT (U/L)* |  |  |  |  |
| 2 h | 43.92±2.311 | 71.43±1.792* | 61.25±2.766*.<sup>#</sup> | 52.74±2.389<sup>#</sup> |
| 4 h | 43.92±2.311 | 81.17±0.943* | 70.23±2.490*.<sup>#</sup> | 60.51±2.006*.<sup>#</sup> |
| 24h | 43.92±2.311 | 63.58±2.588* | 52.95±0.916<sup>#</sup> | 46.81±1.468<sup>#</sup> |
| *AST (U/L)* |  |  |  |  |
| 2 h | 80.51±2.797 | 113.6±2.635* | 96.79±1.961*.<sup>#</sup> | 90.60±2.031<sup>#</sup> |
| 4 h | 80.51±2.797 | 126.3±2.919* | 109±3.220*.<sup>#</sup> | 100.5±3.591*.<sup>#</sup> |
| 24h | 80.51±2.797 | 105.6±4.703* | 89.78±3.644<sup>#</sup> | 84.17±3.017<sup>#</sup> |
| *ALP (IU/L)* |  |  |  |  |
| 2 h | 91.45±2.378 | 129.6±1.651* | 111.4±4.534*.<sup>#</sup> | 100.9±3.855<sup>#</sup> |
| 4 h | 91.45±2.378 | 140.3±1.829* | 120.1±0.779*.<sup>#</sup> | 109.1±2.588*.<sup>#</sup> |
| 24h | 91.45±2.378 | 119.8±4.942* | 104.3±2.045<sup>#</sup> | 95.94±0.676<sup>#</sup> |

\* Significant difference from control group ( $P \leq 0.05$ ).

### 3.2. Histopathology analysis

Microscopic examination of liver from rats after 2, 4 and 24 h of injection with the sublethal dose of native venom (0.672 mg/kg) observed obvious morphological changes in liver tissues. The alterations started with sinusoidal dilatation, congestion, appearance of occasional fat droplets around the dilated central vein after 2 h (Fig. 1 II), then formation of dense fibrous tissue within the portal area and wall thickness of portal vein, bile duct and hepatic artery after 4 h post envenomation (Fig. 1 III), ended with liver cell loss and vacuolated hepatocytes after 24 h of venom injection (Fig. 1 IV). While, the administration of 1.5 kGy and 3 kGy gamma irradiated venom showed less histological alterations than the native one as shown in (Fig 2 V, VI, VII) and (Fig 3 VIII, IX, X), respectively. In addition, the liver tissues appeared almost normal in rats envenomed with the subletal dose of 3 kGy gamma irradiated venom, particularly after 24 h of injection.

**Fig 1.**
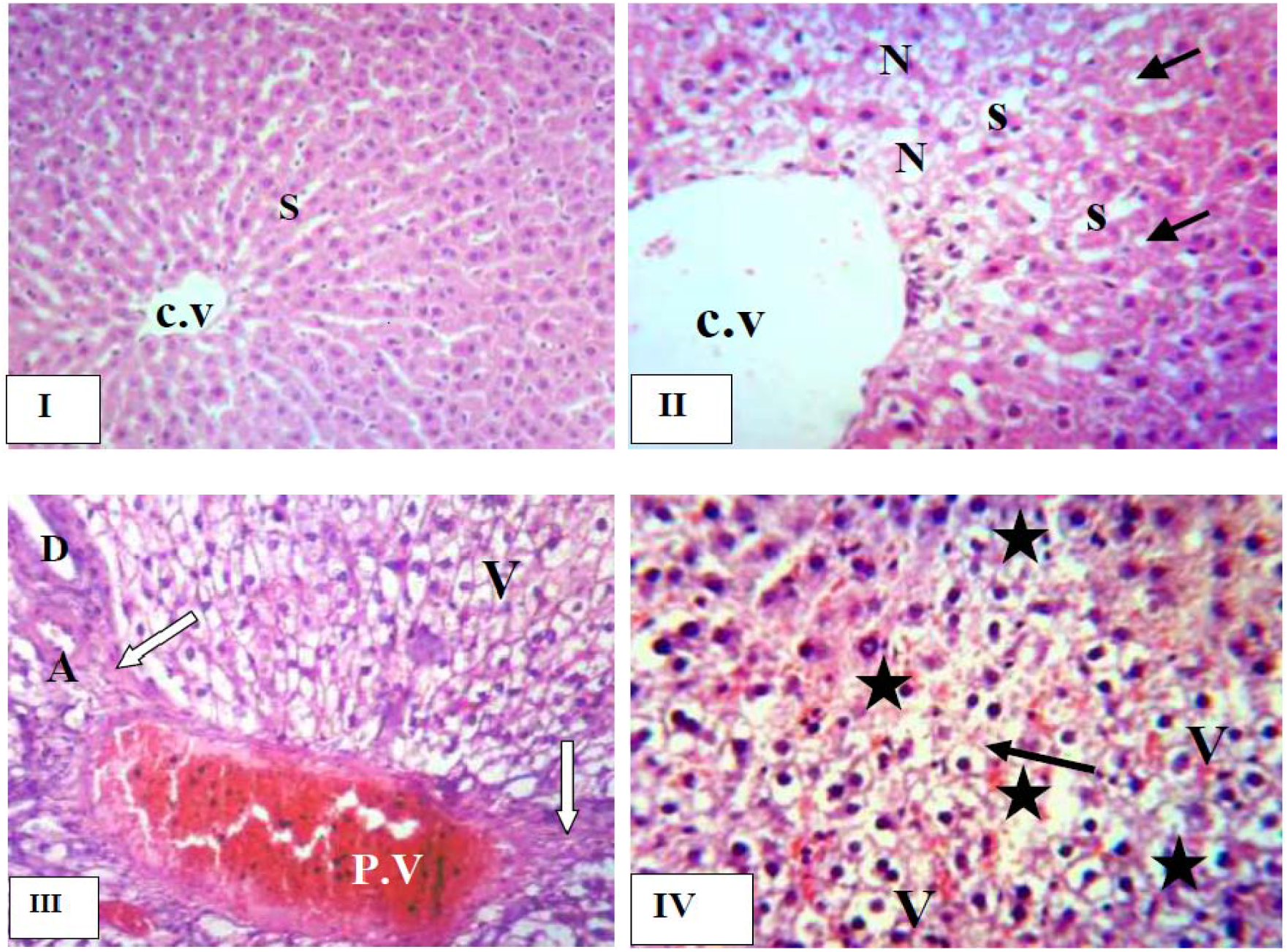
Histological examination of liver of control rats **(I)** a normal architecture of liver tissue with normal hepatocytes, central vein (C.V) and thin sinusoid (S), histopathological alterations of rats’ liver inoculated with native venom showed after 2 h **(II)** sinusoidal dilatation (S), liver cell necrosis (N), occasional fat droplets (arrow) appears around the dilated central vein (C.V), after 4h **(III)** vacuolar degeneration of hepatic cells (V) and enlarged portal tract with dilated congested portal vein (P.V), dense fibrous tissue (arrow) within the portal area and thickened wall of portal vein, bile duct (D) and hepatic artery (A), and 24 h **(IV)** liver cell loss (star), hemorrhage infiltration (arrow) and vacuolated hepatocytes (V) with darkly stained pyknotic nuclei. **(H&E×200).**

**Fig 2.**
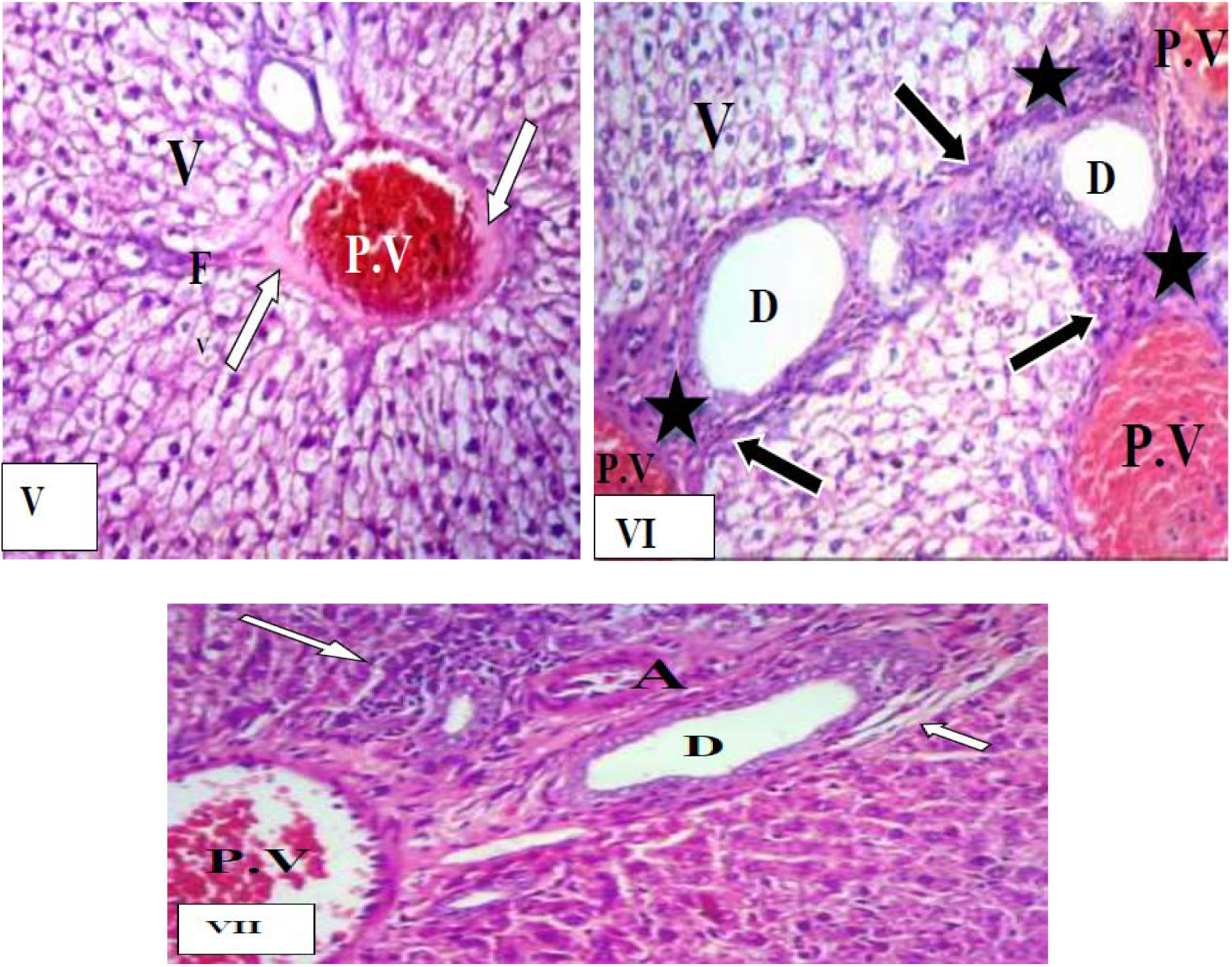
Histopathological alterations of rats’ liver injected with 1.5 kGy gamma irradiated venom showed after 2 h **(V)** vacuolated hepatocytes (V) and congested portal vein (P.V) lined by a layer of dense hyaline (arrow) with thin fibrous tissue (F) radiating from portal tract, after 4 h **(VI)** vacuolar degeneration (V) of hepatocytes, dilated congested portal vein (P.V) with central dilated bile ducts (D) surrounded by inflamed portal tract fibrous tissue, large fibrous septa (arrow) dissecting the parenchyma and linking portal tracts and intense inflammatory cell infiltration (star) of all portal tracts, and after 24 h **(VII)** expanded fibrotic (short arrow) portal area, dilated congested portal vein (P.V), intense inflammatory cells (long arrow) and hepatic artery (A) and bile duct (D) with a thickened wall. **(H&E×200).**

**Fig 3.**
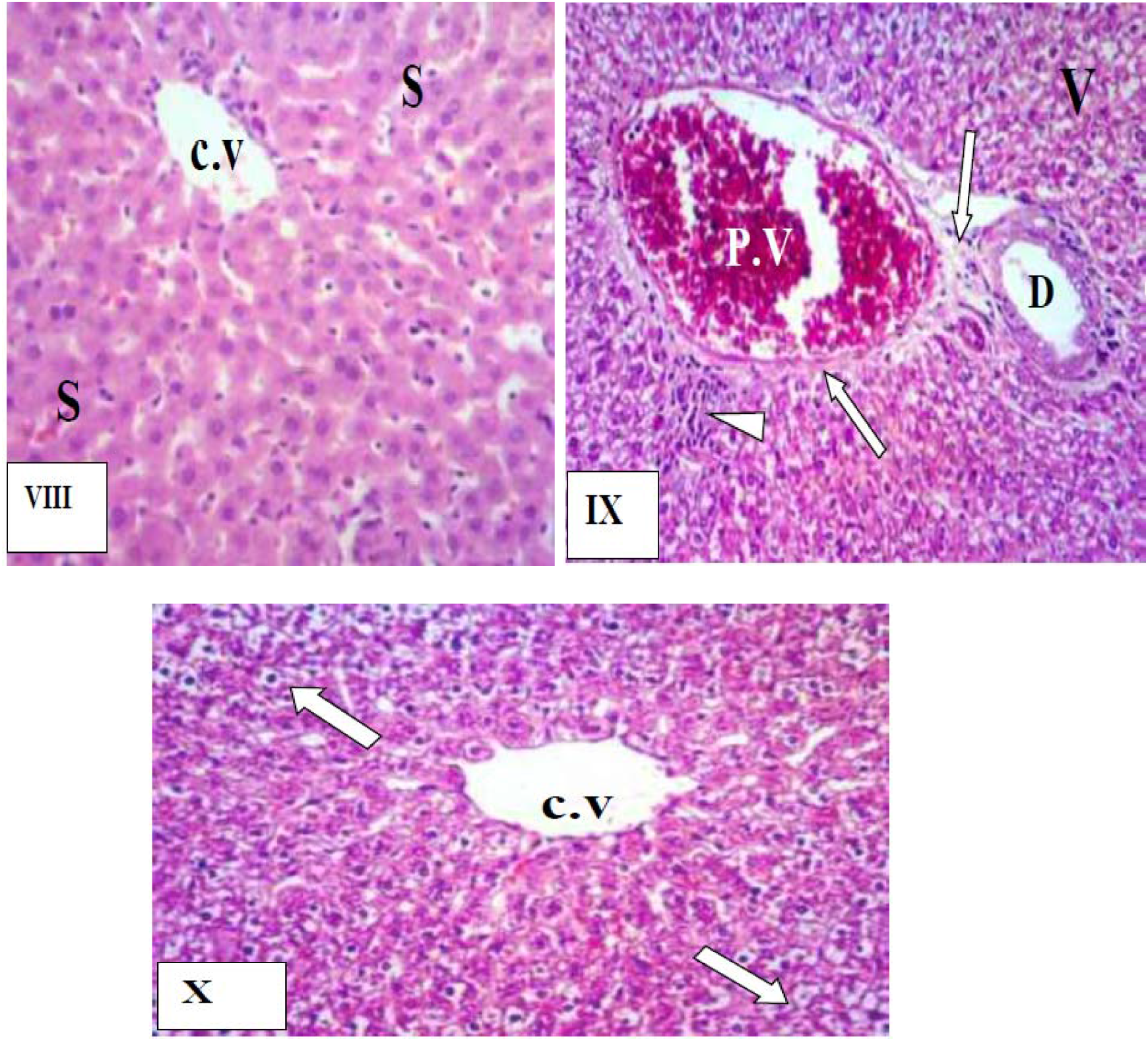
Histopathological alterations of rats’ liver injected with 3 kGy gamma irradiated venom showed after 2 h **(VIII)** normal central vein (C.V) with few inflammatory cells around it, some pericentral sinusoids (S) are infiltrated with red cells; however the remaining liver cells are unaffected and appear normal, after 4 h **(IX)** dilated congested portal vein (P.V) with bile duct proliferation (D), excess fibrous tissue (arrow) and inflammatory cells (head arrow) are present in the portal area, some hepatocytes are vacuolated (V), whilst the remaining liver cells appeared normal and after 24 h **(X)** normal central vein (C.V) with an almost normal appearance of the surrounding liver cells but vacuolation (arrow) of some peripheral hepatocyte is still noted.

## 4. Discussion

*Echis Coloratus* belongs to the *Viperidae* family which comprises *Echis carinatus*, *Echis pyramidum*, *Cerastes cerastes* and *Pseudocerastes fieldi* in Egypt. Its venom induces complex biological effects and multisystem affection that may lead to mortality ***(WHO, 2016)*.**

In the present study, the results are clearly demonstrated that an intraperitonial injection of sublethal dose (0.672 mg/kg) of *Echis Coloratus* venom shows a significant elevation in serum glucose level compared to the control groups after 2, 4 and 24 hours of injection. This is in agreement with ***Salman (2009)*** and ***Jarrar (2010)*** who reported a significant increase in glucose level in the sera of guinea pigs and sheep envenomed with the *Echis Coloratus* venom. This finding may be explained due to several reasons such as activation of β-receptor ***(Corrado et al., 1968)***, release of tissue and medullary catecholamines ***(El-Asmar et al., 1974)***, the presence of target components of the venom themselves could induce hyperglycemia due to inhibition of glucose uptake by skeletal muscle and inhibition of insulin release inducing by target toxins **(*Ismail and Osman, 1973)***, glucose utilization retardation at the peripheral tissues ***(Mohamed et al., 1965)*** and mobilization of glycogen in the liver which may be due to direct inhibition of glucokinase or indirectly by adrenaline release stimulation ***(Mohammad et al., 1980; Al-Saleh, 2002)*.**

The liver is a major producer of most serum proteins and regulates their total levels in the blood, so the sera ALT, AST and ALP levels are known to be good markers for hepatic dysfunction ***(Zilva and Panmall, 1984; Al-Quraishy et al., 2014)***. The results show a significant increase in these enzyme activities after a single injection of sublethal dose (0.672 mg/kg) of native *Echis Coloratus* venom compared to the control groups after 2, 4 and 24 h. These elevations give evidence about the destruction of the liver tissues and liver inflammation as a result of venom injection. This is in accordance with several studies as ***Al-Jammaz, (2003)*** who reported that the activities of ALT, ALP and AST were significantly increased in the sera of rats after 4, 8, 12, 24, 48 and 72 h after envenomation. In addition, ***Jarrar (2010)*** showed a significant increase of these enzymes after *Echis Coloratus* venom administration. Moreover, these results was confirmed by the finding of ***Al-Sadoon and Fahim (2012)*** as they reported a time dependent increase in the levels of AST, ALT and ALP after an intraperitonial injection of a dose equal to LD_50_ of *Echis Coloratus* venom. Accordingly, ***Omran and Abdel Rahman (1992)***, and ***Shaaban and Hafez (2003*)** as they reported that lethal and sublethal doses of venom were capable of triggering stress reactions and increased levels of the ALT, AST and ALP in blood circulation leading to severe damage in many vital organs including the liver.

These biochemical results are in harmony with the histopathological findings of the liver. The *Echis Coloratus* venom without radiation exposure induces loss of the common architecture of liver at the experimental rats after 2, 4 and 24 h of envenomation. Several histopathological manifestations such as cytoplasmic vacuolization, cell swelling, dilation of blood vessels, hemorrhage and inflammation indicate the toxic effects of *Echis Coloratus* snake venom as well as it affects the functions of the liver severely. These findings corroborate with those described by ***Jarrar (2010)*** who observed many histological alterations include cytoplasmic vacuolation, necrosis, hepatocytes atrophy, sinusoidal dilatation, hepatic architecture distortion, central vein congestion and the severely affected hepatocytes appeared swollen with small lipid droplets in the liver of sheep following *Echis Coloratus* envenomation. Additionally, the progressive dilatation of blood vessels may be considered as a reactive change related to an inhibitory effect on vascular smooth muscles which cause relaxation and consequent vasodilatation ***(Jeremy et al. 1990)*.** Meanwhile, the observed inflammatory cellular infiltration may be a secondary effect to the engorgement of blood sinusoids ***(Mikhalidis and Dandona, 1990)*.** Nuclear alterations mainly pyknosis induced by *Echis Coloratus* venom might be due to increased cellular activity and nuclear interruption associated with venom detoxification. The liver necrosis may be explained as a result of the action of venom phospholipase A_2_ enzymes that hydrolyze phospholipids in the cell membrane leading to rupture of the hepatic cell membranes and cellular damage as seen in the present study. The appearance of lipid vacuoles within the hepatocytes might indicate venom interference with mitochondrial and microsomal function that leads to disruption of lipoprotein and resultant lipids accumulation ***(Jarrar, 2010)***.

As venom contains a variation of enzymes such as proteases, phospholipases A2, hyaluronidases, L- aminoacid oxidases, phosphodiesterase and other enzymes that may increase generation of oxidative promoters (hydrogen peroxide, free radicals and lipid peroxides) and decrease the enzymatic activities of antioxidant enzymes (glutathione reductase, superoxide dismutase) leading to induction of oxidative stress and activating key cellular hallmark events including mitochondrial alterations and DNA damage, thus triggering apoptosis ***(Ghneim, 2017)*.**

As regard to the radiation effect on biochemical parameters, a single injection of sublethal dose (0.672 mg/kg) of 1.5 kGy gamma irradiated venom showed significant elevation in ALT, AST, ALP and glucose activity compared to the control group after 2 and 4 hours. Whereas, it did not cause any significant elevation in these parameters activity compared to the normal groups after 24 h. Moreover, a single injection of sublethal dose (0.672 mg/kg) of 3 kGy gamma irradiated venom did not cause significant elevation in rats serum activities of ALT, AST, ALP and glucose compared to the control group after 2 and 24 hours. Although, it showed significant elevation in serum levels of these parameters after 4 h compared to the control group. Beside the biochemical results, injection of 1.5 kGy and 3 kGy gamma irradiated venom showed less alterations than the native one or mostly normal hepatic tract after 3 kGy gamma irradiated venom envenomation. In the same line ***Abib and Laraba-Djebari (2003)*** reported that no tissue alterations were apparent in the myocardium of mice injected with 2 kGy irradiated venom. The disorganization of the molecular structure of venom proteins after exposure to gamma radiation may be explained due to a change in a critical side chain or a break in the hydrogen or disulfide bonds which result in a disorganization of the internal relationships of side chain groups or an exposure of amino-acid groups, leads to a change in toxicity and enzymatic activity of venoms ***(Shaaban et al., 1996; Hayes and Kruger, 2001)*.**

## 5. Conclusion

On the light of the above study, the low doses of gamma radiation is considered as an effective venom detoxification tool helps to reduce the venom toxicity and extends the lifespan of the hyperimmunized animals to decrease antivenin production cost.

## Acknowledgement

The author would like to thank ***Prof. Dr. Manar N. Hafez***, Professor of Histology, Radiation Biology Department, National Center for Radiation Research and Technology, Atomic Energy Authority for her valuable help in reading and confirming the histopathological alterations.

## Ethical approval

All animal handling procedures were approved by the Ethics and Animal Care Committee at Faculty of Pharmacy, Cairo University which obeys the National Institutes of Health guide for the care and use of Laboratory animals (NIH Publications No. 8023, revised 1978).

